# KCFtools: Rapid alignment-free method for introgression screening and GWAS using k-mer profiles

**DOI:** 10.1101/2025.11.01.685998

**Authors:** Sivasubramani Selvanayagam, Jesus Quiroz-Chavez, Ricardo H. Ramirez-Gonzalez, Cristobal Uauy, Sandra Smit, M. Eric Schranz

## Abstract

**Motivation:** In the era of multiple genome references, researchers often align sequencing reads against distinct assemblies or even multiple references simultaneously. This enables applications such as the detection of introgressed segments or highly variable genomic regions, which are especially prevalent in large-genome crop species such as lettuce or wheat. However, these applications come at the cost of increased computational burden, inconsistencies in mapping methods, and reduced reproducibility across studies. To address these limitations, we developed KCFtools, a Java-based toolkit that identifies the presence and absence of *k*-mers in non-overlapping genomic or transcriptomic windows by comparing query and reference genomes. This alignment-free approach enables the efficient computation of an identity score for each window, thereby facilitating robust detection of introgressed or variable regions across genomes.

**Results:** We systematically evaluated the performance and accuracy of the *k*-mer-based method implemented in KCFtools, benchmarking it against conventional SNP-based introgression detection pipelines. Our results demonstrate that KCFtools effectively captures introgressed segments and structurally diverse regions, even in species with fragmented or highly divergent reference genomes. In addition, we extended KCFtools to generate genotype matrices from *k*-mer variation tables. These matrices are compatible with Genome-Wide Association Studies (GWAS) software and allow the identification of loci associated with phenotypic traits. We showcase the utility of this approach by detecting known and novel associations for downy mildew resistance in lettuce, underscoring the pipeline’s potential for high-resolution, reference-agnostic population genetic analysis.

**Availability:** https://github.com/sivasubramanics/kcftools

## Introduction

Understanding and harnessing genetic diversity is important to both evolutionary biology and modern crop improvement (Bohra et al., 2022; Swarup et al., 2021). Introgression, the transfer of haplotypes between species or populations through hybridization and backcrossing, has historically been essential in domestication and adaptation by introducing beneficial traits such as stress tolerance and disease resistance (Hufford et al., 2012; Tiffin and Ross-Ibarra, 2014). Recent advancements in technologies and the development of chromosome-scale reference assemblies have enhanced our ability to explore genomes in detail, even in species with large and complex genomes like wheat and oat (Krattinger and Keller, 2022; Zhou et al., 2020). The shift towards extensive large-scale population re-sequencing and pangenomic frameworks has uncovered structural and Presence/Absence Variation (PAV) that contribute to agronomic traits (Gao et al., 2019; Jayakodi et al., 2021). Nonetheless, the comprehensive use of these genetic resources for identifying introgression and variable regions requires robust computational approaches capable of handling the scale, complexity, and diversity of genomics datasets.

Introgression can have diverse evolutionary and agronomic importance, ranging from adaptive gains to neutral or even maladaptive effects. Adaptive introgression introduces alleles that enhance fitness traits such as stress resilience or pathogen resistance, and these variants are often retained through selection (Hedrick, 2013; Whitney et al., 2006). In contrast, neutral introgression contributes to background genetic diversity, while maladaptive introgression may reduce local adaptation or compromise fitness (Hufford et al., 2012). In crop species, adaptive introgression from wild relatives has been instrumental in improving traits related to productivity and environmental tolerance. For example, in lettuce (*Lactuca sativa*), genomic analyses have revealed introgressed regions from wild progenitors such as *Lactuca serriola, Lactuca virosa*, and *Lactuca saligna* that contribute to disease resistance and root architecture (Parra et al., 2021; Uwimana et al., 2012), in wheat, over half of the cloned disease resistance genes originate from wild wheat species (Hafeez et al., 2021). Such cases illustrate the potential of introgression to enrich genetic diversity and enhance agronomic performance, underscoring the need for precise methods to identify and characterize these genomic regions. Tools such as GWAS and genetic mapping are critical to harness such regions by linking them to phenotypic variation and guiding their use in breeding programs.

Identification and characterization of introgressed and/or variable regions are essential for both fundamental research and applied breeding, as they allow researchers to discover genomic segments contributing to adaptive and/or agronomic traits introduced from wild relatives (Hufford et al., 2012). However, these methods can be expensive in species with large and complex genomes, such as polyploids or those with high repetitive content, where read mapping and variant calling become unreliable (Edelman et al., 2019). The computational burden is further compounded by the increasing number of query genome references, as traditional introgression detection methods require pairwise comparisons that scale poorly with the size of genome size and datasets. Furthermore, distinguishing true introgression from incomplete lineage sorting remains a major challenge in alignment-dependent frameworks (Martin and Jiggins, 2017), resulting in possible misidentification or undetected adaptive loci.

Traditional approaches for detecting introgression generally rely on Single Nucleotide Variation (SNV)-based linkage disequilibrium patterns, phylogenetic inference, D-statistics and Identical By State (IBS) (Aflitos et al., 2015; Patterson et al., 2012). Recent advances, such as haplotype-based statistics or model-based likelihood inference, provide improved sensitivity or specificity in certain contexts (Koppetsch et al., 2024; Hibbins and Hahn, 2022). Nevertheless, these approaches are frequently limited by computational expense, reduced power in complex or polyploid genomes, and an inability to confidently distinguish introgression from similar signatures (e.g., incomplete lineage sorting). This has prompted the need for more flexible, robust, high-resolution tools to explore genome variation beyond the limitations of conventional mapping-based analyses.

Recent advances in pangenomics have revolutionised our understanding of crop genome diversity by capturing both core and dispensable genes across populations, enabling a broader context for detecting structural variants and introgressions (Gao et al., 2019). Pangenome graphs and assemblies facilitate the identification of PAV and novel sequences that are otherwise missed in single-reference studies (Jayakodi et al., 2021). Supporting that *k* -mer-based methods have been applied in various such genomic analyses, including genome size estimation and detection of structural variants (Sun et al., 2018). However, there remains a lack of dedicated tools that leverage *k*-mer-based approaches specifically for genome-wide identification of introgressions.

Here, we present the KCFtools pipeline and its applications in detecting introgressions and variable regions between the reference genome and query genomes, as well as sequenced data, by advancing the *k* -mer based method used to detect introgression(Ahmed et al., 2023). We comprehensively evaluate the performance and accuracy of this *k* -mer-based method, benchmarking it against the conventional SNV-based introgression detection techniques. Further, we extend the functionality of KCFtools in generating genotype matrices from *k* -mer variation tables. These genotype matrices can be used in GWAS, leveraging the power of this pipeline for screening genetic variants associated with phenotypic traits of interest. Additionally, we have implemented functionalities to handle RNAseq data to screen for introgression from transcriptome data.

## Methods

The KCFtools pipeline facilitates the detection and characterization of genomic variation at the *k* -mer level across diverse genotypes. KCFtools supports a range of applications based on window-based identity scoring, adaptable to different sequencing data types and research objectives:

- **Introgression and Variable Region Detection:** Using sequenced reads or whole genome from introgressed genotypes and a reference genome (donor or recurrent), KCFtools identifies genomic windows with significantly reduced identity, indicative of introgressed or highly variable regions.
- **Genotype Matrix Construction for GWAS:** From multi-sample sequencing data, KCFtools computes identity-based genotype matrices, enabling genome-wide association studies, kinship estimation, and population structure analysis.
- **Transcript-Level Variation from RNA-seq:** By analysing RNA-seq reads querying against the annotated reference genomes, KCFtools detects gene/transcript-level presence/absence variation, facilitating expression-informed introgression mapping and gene PAV discovery.

An overview of the KCFtools workflow, outlining the core processing steps and their integration across the different applications described above, is presented in Figure 1. This pipeline structure serves as a common framework, providing a unified basis for introgression detection, transcript variation analysis, and genotype matrix generation.

**Fig. 1:**
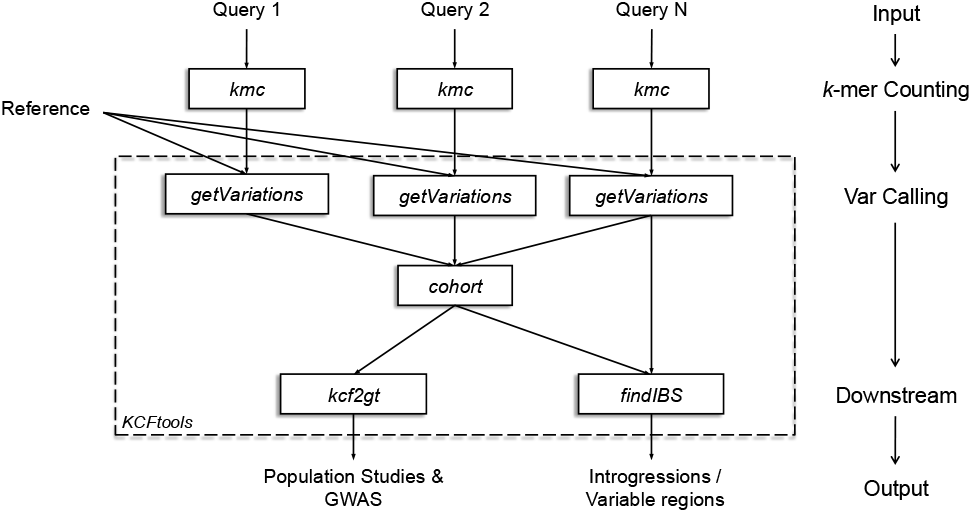
Schematic overview of the KCFtools pipeline. The processing steps involving external tools and plugins are denoted in italics, while the plugins that are part of KCFtools are denoted within the dotted box. The cohort step can be used only when two or more queries are used in the pipeline, enabling the comparative analysis. The reference data mentioned here is in multi-fasta format (e.g., chromosomes), and the query data could be in multi-fasta/fastq format.

### Variation Screening

To detect *k* -mer PAV across genomes, we implemented a Java-based framework inspired by Identity-by-State in Python (IBSpy) (Ahmed et al., 2023) leveraging *k* -mer counting via KMC3 (Kokot et al., 2017). The method focuses on splitting the reference genome into non-overlapping (or sliding) windows and assessing the presence of *k* -mers derived from the reference in query sequences. The *k* -mer table for the query is generated using KMC3, a highly efficient *k* -mer counting tool optimized for large-scale genomic datasets, producing a signature indexed hash table. These reference windows can be simple, continuous, non-overlapping genomic segments; transcript sequences derived from GTF features (for RNA-seq data); or gene sequences, enabling gene presence-absence variations screening at different resolutions. For each window, the number of observed *k* -mers is counted, along with the number of variations (gaps) between consecutive *k* -mer hits. The *k* -mer distance is calculated as the number of consecutive bases that are not covered by the observed *k* -mers. This *k* -mer distance serves as a measure of sequence divergence within the specified window. The overall *k* -mer distance is partitioned into two components (Supplementary Figure S4): the ‘inner distance’, representing the aggregated gap lengths flanked by observed *k* -mers within the window, and the ‘tail distance’, which quantifies the cumulative gap lengths at either end of the window, effectively capturing the extent of *k* -mer absence at the window’s boundaries. The overview of this methodology is presented in Figure 2.

**Fig. 2:**
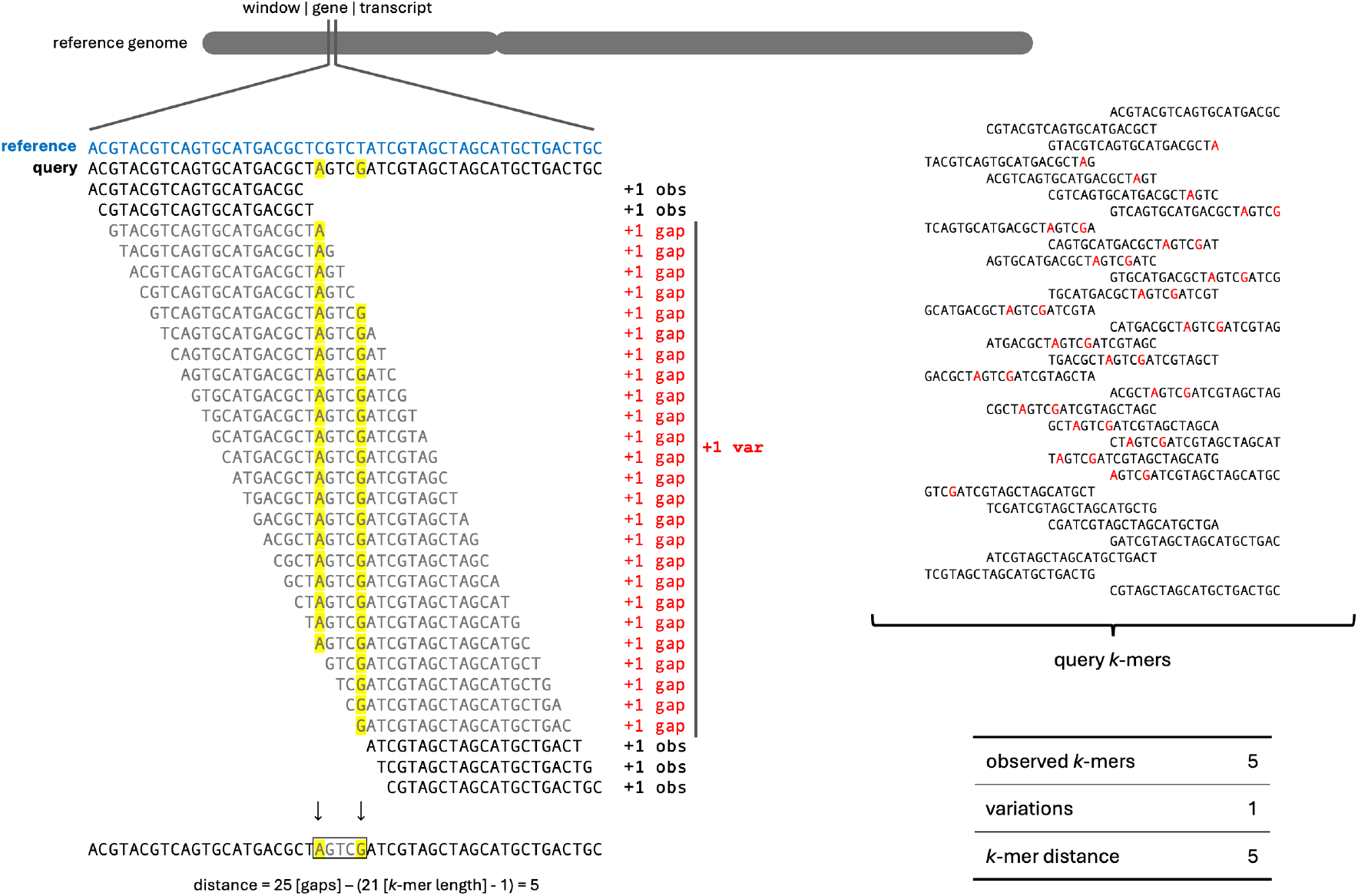
Schematic overview of the methodology of KCFtools and how the *k* -mer-based variation attributes are calculated between the reference genome and the query sequences.

### Scoring windows

To quantify the overall sequence similarity between reference and query within each window, we calculate an Identity Score (*IS*) based on *k* -mer presence and the distribution of *k* -mer distances. The score is a weighted sum of three components, according to the following formula:

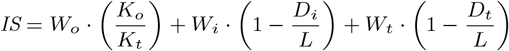

Where *K*_*o*_ is the number of observed *k* -mers in query *k* -mer db, *K*_*t*_ is the total number of *k* -mers in the reference window, *D*_*i*_ is the inner distance, *D*_*t*_ is the tail distance, and *W*_*o*_, *W*_*i*_, *W*_*t*_ represent the weights assigned to the *k* -mer ratio, inner distance ratio, and tail distance ratio, respectively. The effective length (*L*) represents the overall *k* -mers covering length of the window (i.e., window length + *k* -mer size - 1). This Identity Score provides a comprehensive measure of sequence identity, considering both the density of observed *k* -mers and the distribution of gaps within the window.

### Introgression vs Variable windows

Depending on the choice of reference and query used to calculate the *k* -mer variations, we could classify genomic regions as possible introgressions or variable regions in the query data (Supplementary Figure S7). We used an identity score threshold to distinguish these introgression and variable regions, as consecutive genomic windows must meet the defined identity score cutoff to be classified as introgressed or variable windows. In case the introgressed query is compared against a reference genome representing the donor genotype, consecutive windows exceeding the identity score threshold are classified as introgressions, indicating the segments of the donor genome present within the query. Alternatively, if the reference genome is of the recipient genotype, windows below the cutoff are classified as variable regions, reflecting genomic divergence between the reference and the query. This classification allows for the identification of conserved and divergent genomic segments, enhancing the detection of variations and introgressed regions across the genome.

### *k* -mer variations as Genotype Matrix

Building on the methodology described, we extended our approach to screen for *k* -mer variations across multiple genotypes using the developed framework. By employing the cohort plugin, we integrated *k* -mer variation data from multiple query samples into a consolidated KCF file, which efficiently stores the presence/absence matrices of *k* -mers across multiple samples. Based on identity score thresholds, the consolidated KCF file can be transposed into a genotype matrix format analogous to a traditional germline genotype matrix. In this representation, each data point is encoded as follows: 0 for homozygous reference alleles, 2 for homozygous alternative alleles, and 1 for heterozygous alleles based on the identity score for each window (Supplementary Figure S5 and Supplementary Table S1). This transformation facilitates the interpretation of presence/absence patterns in a genotype-like context, aligning with standard practices in population genomics. The resulting genotype matrix offers a versatile foundation for downstream population genetic analyses. For instance, it can be directly utilized in studies investigating admixture, kinship, and Linkage Disequilibrium (LD). Additionally, this matrix can serve as a genotype matrix for GWAS to identify loci associated with phenotypic traits. The *k* -mer-based genotype matrix approach thus bridges sequence-level variation detection with higher-order genomic analysis, providing a robust framework for exploring genetic diversity and population structure.

### Software architecture

The core pipeline of the KCFtools is a combination of custom plugins written in Java and a set of helper scripts written in Python and R. The core functionalities of the KCFtools package have been structured within a modular, plugin-based architecture (e.g. `findIBS` plugin designed to identify genomic variants), that offers a unified API, enabling both command-line execution and seamless integration with third-party bioinformatics pipelines. The code is freely available, open source, and distributed as a JAR executable. The pipeline plugins, individual components, and example datasets with parameters are all documented in the GitHub repository (https://github.com/sivasubramanics/kcftools).

### Application Testing

A custom python script was written to simulate the introgressions by introducing a variable number of segments from the donor genome by replacing the segment in the recipient genome. We used chromosome 8 (25.47 Mb) of *Oryza glaberrima* as the donor genome, while chromosome 8 (28.6 Mb) of *Oryza sativa* Japonica group as the recurrent genome. `randomreads.sh` from `bbmap` (Bushnell, 2014) package was used to simulate paired-end reads from these simulated chromosomes for 10x depth with snprate 0.5% and indel rate of 0.1%. Selection of *k* -mer size was generic for this dataset, and we have used k=41 and a window length of 5 kb for this exercise, and the pipeline could detect the introgressions with a few kilobases resolution. To evaluate the accuracy of the method, we compared the simulated data for observed and expected introgression lengths for each iteration. For the accuray F-score, recall, and precision were used as metrics. In order to facilitate the comparative read depth analysis we used the same iteration with different depths to simulate the sequencing reads.

To evaluate the applicability of the pipeline in real world data, we analyzed re-sequencing data from 198 *L. sativa* accessions (van Workum et al., 2024). The *k* -mer variation detection was conducted using the wild genome of *L. virosa* (Xiong et al., 2023) as the donor genome and *L. sativa* (Reyes-Chin-Wo et al., 2017) as the recipient genome to identify introgressed and variable genomic regions. Following this, we transposed the variable windows to a genotype matrix, which was subsequently utilized for population studies and GWAS. Kinship and GWAS analysis were performed using GAPIT (Lipka et al., 2012). To validate the accuracy and efficacy of our *k* -mer-based GWAS approach, we compared the identified significant signals with those reported in the previous study by Wei et al. (2021).

## Results

### Features and Core Capabilities

KCFtools introduces a comprehensive approach to detect introgressions using a *k* -mer-based methodology, optimized for comparative genomics and downstream genotype matrix construction. The framework is composed of a suite of Java-based plugins supported by R scripts, enabling integration into both standalone and pipeline-based workflows. The methodology begins with efficient *k* -mer counting using the external tool KMC3, followed by variation screening through window-wise identity scoring. The `findIBS` plugin allows for the detection of introgressed and variable regions based on identity scores, while the cohort plugin supports cross-genotype comparisons by merging individual KCF profiles into a unified KCF file. This facilitates the transformation of window-wise identity scores into a genotype matrix compatible with tools for population structure analysis and GWAS. KCFtools supports *k* -mer count table generated using `kmc` and genome multi-fasta and can operate at multiple genomic resolutions, including fixed-size non-overlapping windows and gene models from GTF annotations. This allows the framework to be tailored to specific biological questions and dataset types (e.g., DNA-seq, RNA-seq).

### Introgression and Variable Regions on Simulated Data

To validate the accuracy of the KCFtools method, we simulated introgressions on one of the rice chromosomes between two different species. The impact of introducing error rate in the reads simulation has been witnessed in the `getVariations`, due to which a few of the false-positive variable windows were observed during the `findIBS` step (Supplementary Figure S9.b and S9.c). These short variable windows could be filtered out based on the “IBSproportion”, which is defined as the proportion of the IBS window blocks against the total windows in the region. Further each iteration were evaluated with observed vs expected introgression and/or variable region signals and the F-scores were above 0.99 for majority of the iterations.

### Impact of sequencing depth

Just like any methodology, various aspects can impact *k* -mer profiles while working with sequencing reads, a few to mention: type of platform, read quality, read length, and depth. We used simulated iterations at different depths to evaluate the impact of sequencing depth on the *k* -mer profile (k=41) of raw reads. To evaluate the impact of sequence coverage and depths, we analyzed simulated datasets across a range of coverages. Our results (Supplementary Figure S6) indicate that, although variations are observed at lower depths, the *k* -mer ratio becomes stabilized and consistently uniform when sequencing depths are more than 8x for short reads. This finding indicates that higher depths provide enough redundancy to overcome stochastic sampling biases in *k* -mer-based coverage estimation. Further to add, with the lower sequencing depth data, the variation observed could be impacted due to the absence of *k* -mer due to the sequencing depth over the variable signals against the reference.

### Impact of *k* -mer size on runtime

The effect of *k* -mer size on processing time is shown in Figure When using KMC for *k* -mer counting, longer *k* -mers may result in the formation of longer super-*k* -mers array in KMC database, potentially reducing upstream processing time due to fewer redundant operations. Nevertheless, increasing *k* -mer size may negatively impact the runtime of the `getVariations` plugin (Figure 3), primarily due to the additional computational requirement of the inherent search and count operations. In order to compensate for differences in processing time, the plugin maintains a constant signature length of 9, thereby providing consistency in performance. Variable prefix lengths, which are integral in KMC’s `LUT_PREFIX_ARRAY` structure, can have a major influence on runtime. For each *k* -mer size, KMC alternates between prefix lengths of 5 and 3, which can lead to additional inconsistencies in `getVariations` performance. Besides the control provided by the CPU, runtime also depends on memory management. The plugin provides two modes: if the `-m/--memory` option is supplied, the complete *k* -mer table is pre-loaded in RAM, whereas in its absence, the suffix array is memory-mapped, which typically results in comparatively slower performance. For small-to moderate-sized data, enabling the `-m/--memory` option can make a major contribution towards runtime efficiency.

**Fig. 3:**
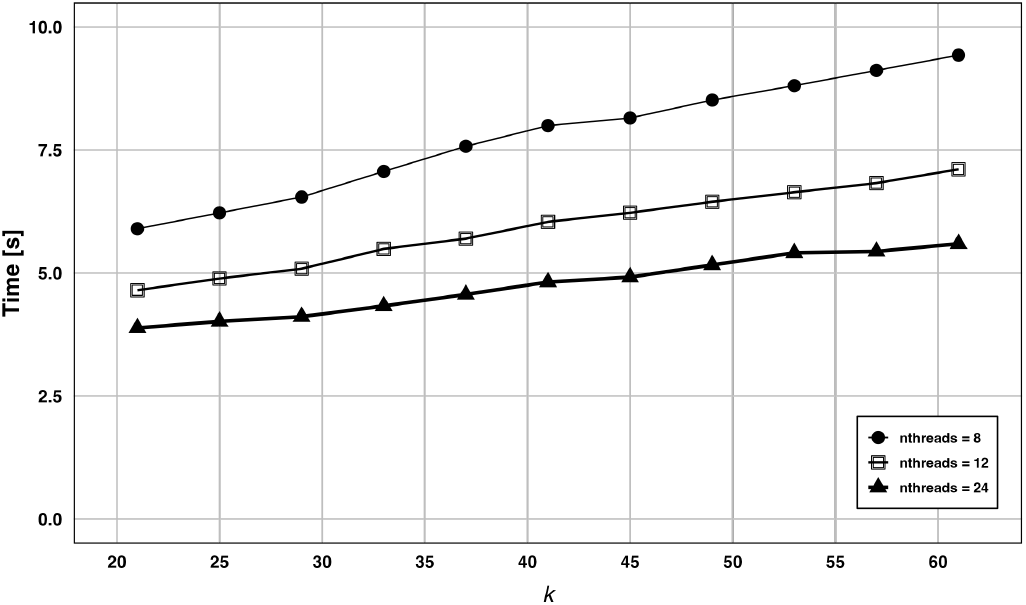
Mean runtime of the KCFtools `getVariations` plugin on simulated data using different numbers of CPU threads (with the `-m` option).

### Impact of feature type on runtime

To obtain a comparative overview of the runtime performance of KCFtools, we analyzed the 198 *L. sativa* WGRS datasets from (van Workum et al., 2024) against the *L. sativa* reference genome (2.4 Gb). Two types of query segments were tested: (i) annotated transcripts and (ii) fixed-size genomic windows. For benchmarking, we also compared these results against a conventional read-mapping and variant-calling pipeline (bwa-mem + deepvariant). The mean runtime were ≈ 8 minutes when using transcripts, ≈ 16 minutes when using genomic windows, and 680 minutes for the conventional pipeline, highlighting the substantial runtime advantage of KCFtools (Supplementary Figure S10). These analyses were conducted on datasets with sequencing coverage ranging from 14x to 40x (mean 22x), ensuring that the performance assessment reflects typical variation in sequencing depth.

### Application 1: Wild introgression in cultivated lettuce

In this exercise, we hypothesized that *L. sativa* (Crisphead-type) accessions may harbor introgressed genomic regions originating from wild *L. virosa*, particularly regions associated with root development and resistance to biotic or abiotic stresses. To investigate this, we employed the KCFtools pipeline to scan for potential introgressions, using the recently published *L. virosa* genome as the reference (Xiong et al., 2023). The analysis was conducted with a *k* -mer size of 31 and a window length of 50 kb, enabling high-resolution detection of candidate introgression events. A prominent introgression signal spanning approximately 25 Mb was identified on scaffold 5 of *L. virosa*, ranging from 125 Mb to 150 Mb (Figure 4). This large segment appears to have been introgressed into a Batavia-type cultivar (LK085 in (van Workum et al., 2024)), suggesting historical breeding or hybridization events between *L. sativa* and *L. virosa*. The presence of such a substantial region indicates possible selection for beneficial traits, potentially contributing to improved stress tolerance or agronomic performance. Additional functional characterization of genes in this region could identify candidate loci that are linked with these traits as targets for marker-assisted selection in lettuce breeding programs.

**Fig. 4:**
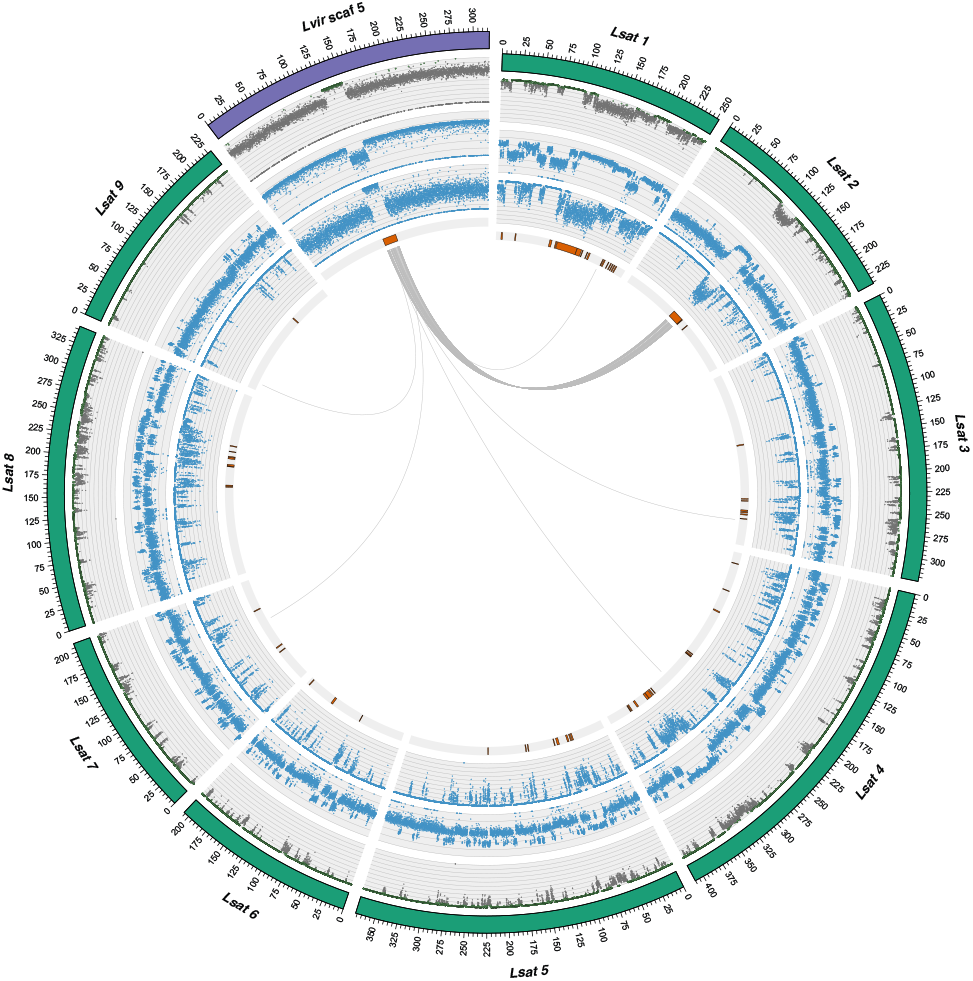
Variable regions in lettuce batavia accession showed against the *L. sativa* genome (green), and the introgression from *L. virosa* scaffold 5 (purple) into *L. sativa* chromosome 2. Rings from outside, a. Chromosome band b. Identity scores c. *k* -mer distance per window (50kb window) d. *k* -mer ratio (observed *k* -mers/total *k* -mers) e. Identified and filtered introgression and variable regions in batavia accession f. Reciprocal Best Hits (RBH) between *L. virosa* and *L. sativa* genes for the target regions

### Application 2: Variable Region screening

Whole-genome re-sequencing data from 198 *L. sativa* accessions, representing a diverse panel commonly used in lettuce breeding and population genetics studies (van Workum et al., 2024), were analyzed to investigate genomic variation across cultivars. Using the *L. sativa* reference genome (Reyes-Chin-Wo et al., 2017), we identified multiple highly divergent genomic regions within the panel (Figure 5.b). Notably, a subset of these regions exhibited population-specific divergence patterns, consistent with signatures of selection in particular cultivar groups. Accurate identification of such regions requires careful optimization of identity score thresholds for each population to mitigate false positives arising from high inter-group variability (Supplementary Information Section 2). When compared with previously reported introgression events (Wei et al., 2021), our findings revealed consistent signals at several major resistance gene clusters, suggesting the persistence of key introgressed loci across breeding cultivars. These conserved regions may reflect targeted selection for disease resistance traits and warrant further investigation to elucidate their functional relevance.

**Fig. 5:**
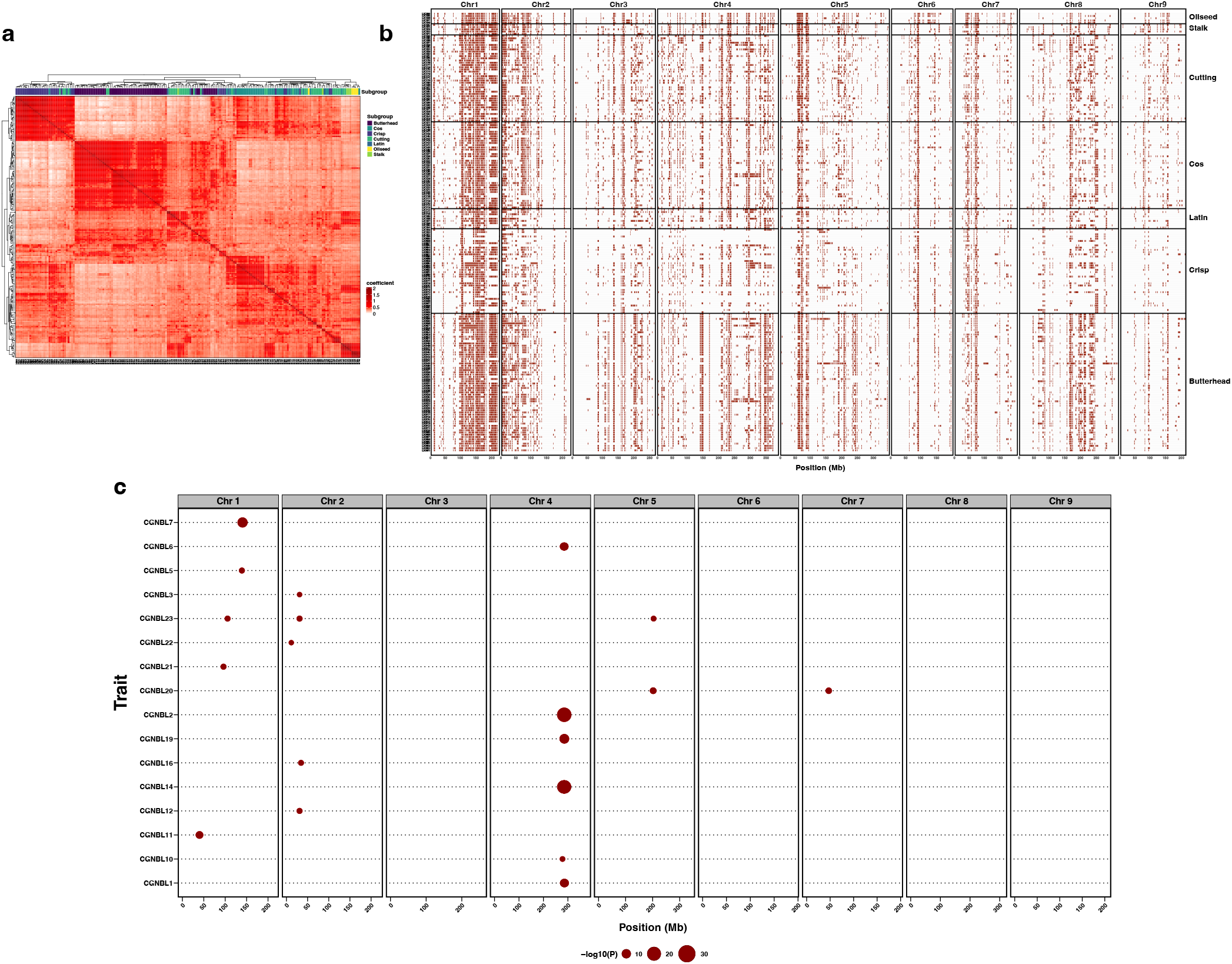
Application of *k* -mer profiles as genotype matrix in *L. sativa* population. a. Kinship plot showing distinct clustering patterns among lettuce accessions, consistent with known subgroups. b. Visualization of variable genomic regions derived from findIBS. c. Manhattan plot from GWAS analysis using the *k* -mer profiled genotype matrix, revealing significant associations for bremia resistance.

### Application 3: Variable regions as genotype data for Population genetics and GWAS

Following identity scoring and introgression mapping, we used the `kcf2gt` plugin to generate genotype matrices from *k* -mer-based identity scores from the results of variable regions screening. Each genomic window was encoded as 0 (homozygous reference), 1 (heterozygous), or 2 (homozygous alternative), facilitating compatibility with GWAS software such as GAPIT (Lipka et al., 2012). The resulting kinship structure (Figure 5.a) revealed distinct clustering patterns corresponding to known subgroups within the diversity panel, reflecting underlying population structure and historical divergence among cultivar types. Further, we performed GWAS with this matrix, combining it with a collection of publicly available *L. sativa* phenotypic datasets, focusing on susceptibility scores to multiple races curated by the Centre for Genetic Resources, the Netherlands (CGN). Resistance to the causal agent of downy mildew, is predominantly mediated by genes encoding Nucleotide-binding Leucine-rich Repeat (NLR) proteins, with occasional contributions from Receptor-Like Kinase (RLK)s and Receptor-Like Protein (RLP)s (Christopoulou et al., 2015b). In lettuce, these resistance genes are often located in major resistance gene clusters that are investigated with the above-mentioned variable regions screening. The GWAS results showcase significant associations localized primarily within a well-characterized resistance gene cluster on chromosome 2 (Figure 5.c), highlighting the utility of KCFtools supported GWAS for uncovering loci associated with pathogen response in lettuce.

## Discussion

Although several *k* -mer-based tools, such as KMC (Kokot et al., 2017), Mash (Ondov et al., 2016), and GenomeScope (Vurture et al., 2017), enable alignment-free analyses of genome composition, they are primarily limited to tasks such as *k* -mer counting, genome size estimation, and similarity assessment. To date, not much of comprehensive framework exists that can robustly detect genomic variation, such as introgressions and/or presence/absence variation, entirely independent of read alignment or conventional SNP-calling pipelines. Recent studies have highlighted the power of *k* -mer–based approaches for population-scale analyses of historical introgression and evolutinaly dynamics (Ahmed et al., 2023; Cheng et al., 2024; Cavalet-Giorsa et al., 2024). KCFtools addresses this methodological gap by leveraging raw *k* -mer frequency profiles to directly identify genomic regions that are polymorphic, construct genotype matrices, and aid in facilitating downstream analyses such as genome-wide association studies. A key strength of KCFtools is its modular, plugin-based design, which separates the pipeline into distinct steps to improve flexibility and maintainability. The findIBS and cohort plugins enable both single-sample and multi-sample analyses, supporting diverse comparative genomics applications. By decoupling the computation of *k* -mer variation from reference genome alignment, the pipeline offers a rapid and alignment-free solution suitable for species with incomplete or divergent reference assemblies.

The application of KCFtools to both simulated and real-world datasets demonstrated its ability to identify introgressed genomic segments with high resolution. The simulated introgressions between *O. glaberrima* and *O. sativa* showcased the ability of the pipeline to resolve even short introgressed segments (5 kb resolution) with high sensitivity and specificity, particularly when sequencing depth exceeds 8×. This suggests that the identity score metric effectively captures both the density and distribution of *k* -mer presence/absence. In the lettuce dataset, KCFtools successfully identified a ≈25 Mb introgressed region from *L. virosa* into cultivated *L. sativa*. The localisation of this region on scaffold 5 suggests targeted selection or historical hybridisation events aimed at introgressing favourable traits such as disease resistance or stress tolerance. The ability of KCFtools to identify such events without requiring a dense SNP map underscores its utility in breeding contexts where non-reference variation plays a crucial role.

A significant implementation of this pipeline is the transformation of window-level *k* -mer identity scores into genotype matrices, enabling integration with GWAS and population studies. The results from our lettuce panel confirm that the *k* -mer-based genotype matrix preserves sufficient resolution and population structure to identify biologically meaningful signals. Importantly, the identification of associations within a known resistance gene cluster (chromosome 4) linked to *Bremia lactucae* resistance validates the biological relevance of the variation (Christopoulou et al., 2015a) captured by KCFtools. The positive overlap between our findings and previous studies further supports the robustness of window-based identity scoring as an informative genomic feature for trait-association studies (Supplementary Figure S8)

Despite its advantages, the performance of KCFtools is limited largely to sequencing depth. We observed that at low coverage (≤8X), stochastic variation in *k* -mer representation can lead to inflated false positive rates in identifying variable regions. Although the IBS proportion metric helps to reduce such noise, applying appropriate coverage thresholds and ensuring consistent sequencing depth across samples remains critical for reliable analysis.

Performance benchmarks demonstrate the impact of parameters such as *k* -mer size, sequencing depth, and memory management on runtime. Longer *k* -mers and higher memory allocations improve specificity but come at the cost of increased computational demands. The availability of the `-m` option and parallelisation support mitigates some of these concerns, making KCFtools suitable for large-scale datasets, provided adequate hardware resources are available. Further to add, we noticed that compared to the conventional mapping-based methods (such as bwa-deepvariant), KCFtools could reduce the analysis runtime by many folds.

## Acknowledgments

We would like to thank Marian Bremer, Marieke Sophia van de Loosdrecht, and Haris Spyridis for their valuable support and insightful discussions regarding the methodology.

## Author Contributions

CU, MES, JQC, and S. Selvanayagam conceptualized the study. RRG, JQC, and S. Selvanayagam developed the software. S. Selvanayagam performed the analysis. S. Selvanayagam, MES, and S. Smit wrote the initial draft. MES and S. Smit supervised the study and performed critical revisions. All authors reviewed and approved the final manuscript.

## Data Availability

The source code and release binaries are hosted at the GitHub repository: https://github.com/sivasubramanics/kcftools. The pipeline script and the scripts used to generate the plots are provided as part of the utility scripts under the same GitHub repository. The Zenodo DOI for the repository: https://doi.org/10.5281/zenodo.15524673.

## Supplementary Information

**Supplementary Material 1:** Supplementary Note: Supporting information about the attributes, definition of terminologies and additional supplementary figures.

**Supplementary Material 2:** Supplementary Table S2) Accession Information used in this study. Supplementary Table S3) Identified variable regions for *L. sativa* accessions. Supplementary Table S4) Kinship matrix for the *L. sativa* accessions. Supplementary Table S5) Runtime information for each accession.

