## Supplementary Material 1 for "KCFtools: Rapid alignment-free method for introgression screening and GWAS using k-mer profiles"

### Supplementary Information for KCFtools

**2 Identity score classification** **4**

**3 Impact of Sequencing Depth** **6**

**4 Additional Supplementary Figures** **8**

### 1 Type of Attributes implemented in KCFtools

#### 1.1 Observed $k$-mers

Based on the reference genome assembly and the $k$-mer database, KCFtools getVariations computes the number of $k$-mers within a given genomic window that are also present in the $k$-mer database of a query sample. The observed $k$-mers metric represents this count. When all $k$-mers in the window are detected in the query sample's $k$-mer database, the observed $k$-mers value equals the total number of $k$-mers in that window. An illustrative example of the observed $k$-mers calculation is provided in supplementary figure S1

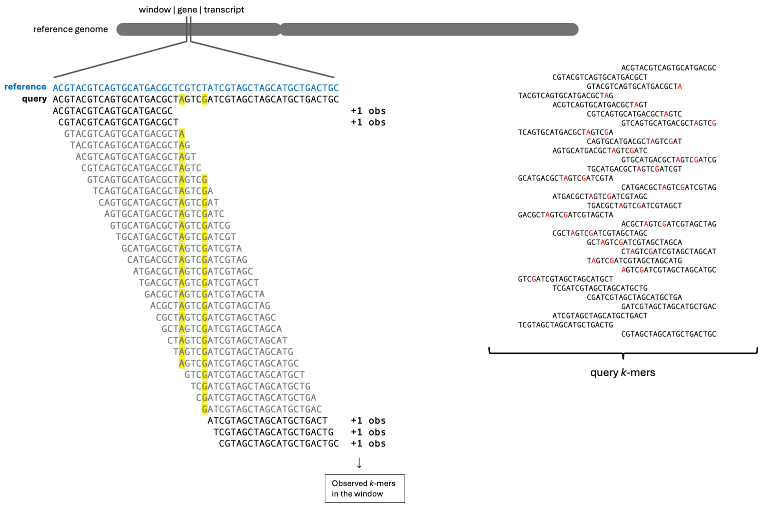

**Supplementary Figure S1: Observed *k*-mers** in window. We compare all the $k$-mers in a reference sequence window to the $k$-mers of a query sample (black) (from raw reads, scaffold-level, or chromosome-scale assemblies) and count the number of observed $k$-mers within each window. The example shows two Single Nucleotide Variation (SNV)s (in yellow) between the reference (blue) and a query (black) and how only the observed $k$-mers present in both the reference and the query are counted.

#### 1.2 Variations

Similar to the observed $k$-mers, KCFtools getVariations plugin scans windows of the reference genome to detect sequence variations. Within each window, the tool examines each $k$-mer sequentially, looking for the presence of it in the query $k$-mer database. If a single $k$-mer or a series of overlapping $k$-mers from the reference are absent in the query sample, this absence is interpreted as a single variation event. This approach allows the detection of both isolated and clustered sequence differences in a high-resolution, $k$-mer-based manner. Two SNPs/InDels whose distance is less than the $k$-mer size used will generate a variation size of '2n-k + the distance' from the first to the last SNV found. In such cases, the tool will record a single variation.

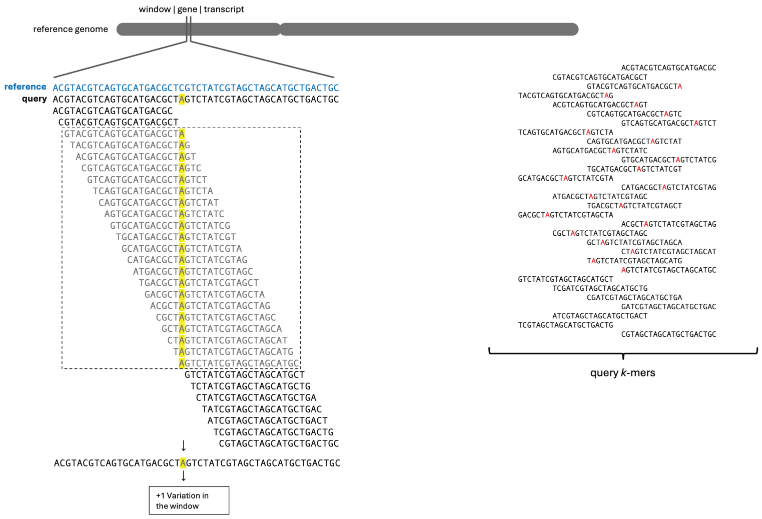

**Supplementary Figure S2: Variations** in window. A variation is a set of continuous overlapping $k$-mers from the reference that are absent in the query. In this example, a single nucleotide difference ($C \rightarrow A$) causes the absence of 21 subsequent $k$-mers.

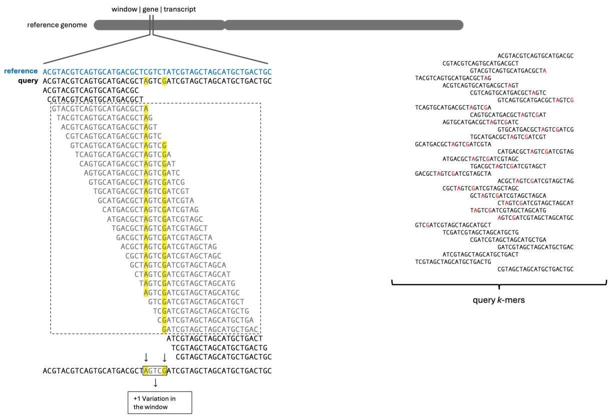

**Supplementary Figure S3: Variations** in window. In this example, multiple single nucleotide differences result in the absence of more than 21 subsequent $k$-mers. Until the next observed $k$-mer is reached, this stretch is considered a single variation.

#### 1.3 $k$-mer distance

When two or more sequence variations (e.g., SNPs or InDels) occur within a distance shorter than the $k$-mer length, they are collectively reported as a single variation. The $k$-mer distance is defined as the number of bases within the window that are not covered or contributed by any observed $k$-mers. Additionally, inner $k$-mer distance refers to the number of uncovered bases that exist between two observed $k$-mers flanking a variation, while tail $k$-mer distance denotes the number of uncovered bases at the trailing end of the window that are not flanked by any observed $k$-mers at either end.

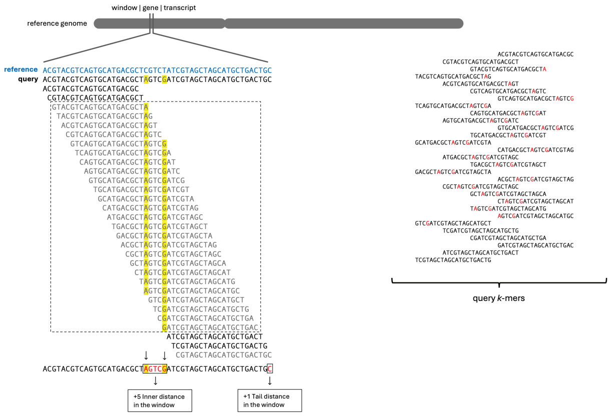

**Supplementary Figure S4: *k*-mer distance** in window. Illustrates the methodology of the inner and tail $k$-mer distance calculations for a window.

### 2 Identity score classification

We hypothesised that sequence-level differences between two samples, such as SNPs or InDels, would be reflected in the differential presence/absence of $k$-mers within defined genomic regions (e.g., windows). By selecting an appropriate $k$-mer size, these local sequence differences can be effectively captured, generating a genome-wide fingerprint of variation across the reference genome. In order to test the hypothesis, we used a Batavia lettuce genotype from [Wei et al., 2021] against the *Lactuca sativa* v11 reference genome. While exploring the variations and identity scores across different chromosomes, we observed that variable pattern of scores defining possible IBS windows (Supplementary Figure S5.b). The next step we intended to define a threshold to categorically assign the windows to different levels of variations (Supplementary Table S1) specific to this data.

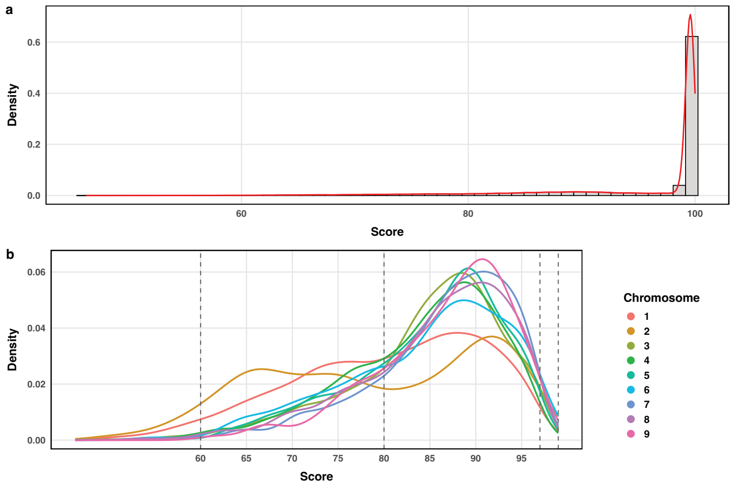

**Supplementary Figure S5:** Identity score distribution of lettuce Batavia against the *L. sativa* reference genome v11 with 50 kb as window size. a. Density plot of the identity score across the whole genome. b. Density plot of per-chromosome identity scores filtered only with less than 97.00 to avoid the skewed distribution.

**Supplementary Table S1:** Interpretation of Score Ranges for Genomic Comparisons

| Score Range (%) | Interpretation | Proposed Category | Biological Implication |
|---|---|---|---|
| >99 | Nearly identical to reference | Reference-like / Unchanged | No significant variation; high-confidence match to the reference genome. |
| 97–99 | Minor variation or heterozygosity | Hybrid-like / Minor Variants | May represent hybrids, residual heterozygosity, or minor SNVs/InDels. |
| 80–97 | Introgression or structural variation | Introgressed / Divergent Regions | Suggests the presence of foreign DNA (e.g., from wild relatives) or large insertions/deletions. |
| 60–80 | Deletions or missing $k$-mers | Putative Deletion / Low Coverage | Likely indicates deletions, or poor sequencing coverage. |
| <60 | Highly divergent or absent | Absent / Homoeologous / Complex | May correspond to homologous regions (e.g., A/B/D genome divergence (Ahmed et al., 2023)) or complex structural rearrangements. |

### 3 Impact of Sequencing Depth

Genome assemblies and $k$-mer based algorithms are usually affected by read length where less coverage is needed with longer reads. Here we tested and validated KCFtools to detect variations using raw reads using different sequence depths and defined an optimal sequence coverage by comparing $k$-mer ratios (observed / total).

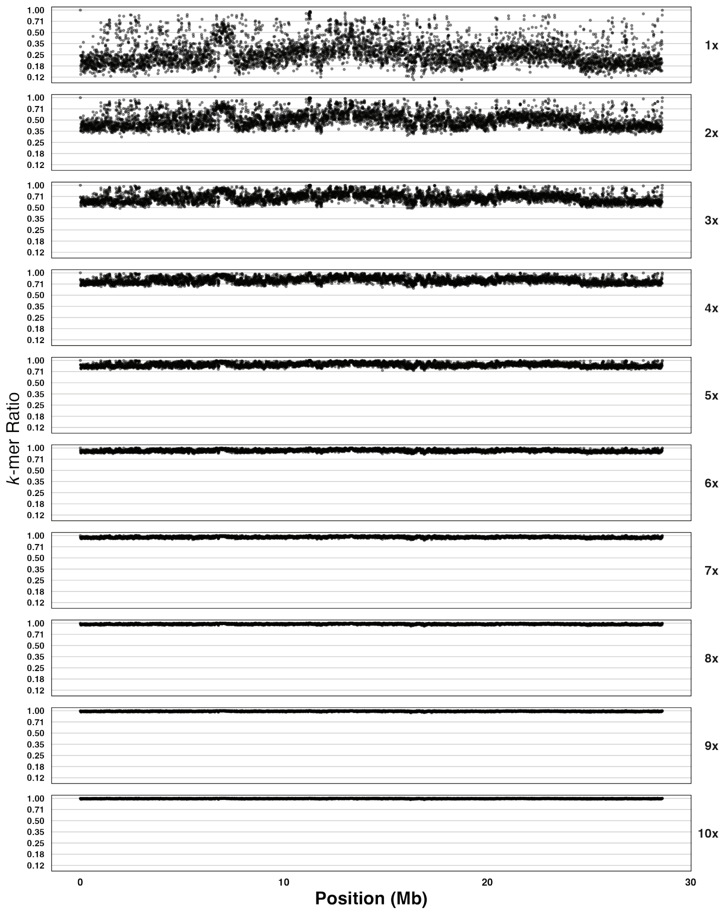

**Supplementary Figure S6:** $k$-mer ratio observed at different sequencing depths using simulated paired-end data with a read length of 100bp per read.

### 4 Additional Supplementary Figures

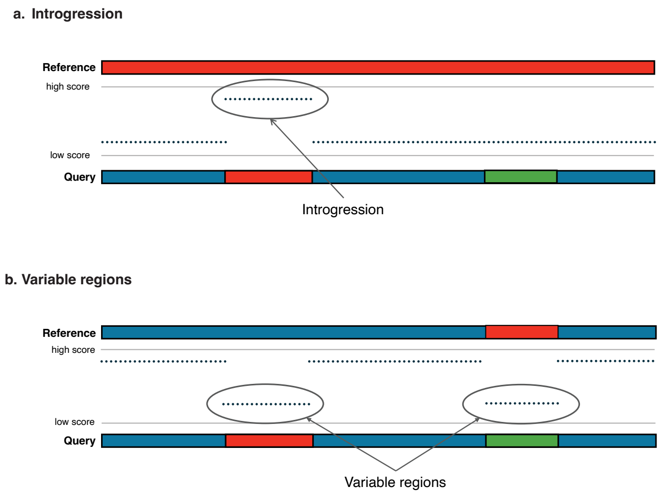

**Supplementary Figure S7:** Schematic of how KCFtools distinguishes (a) introgressed segments and (b) variable genomic regions based on the windowed scores against the chosen reference genome.

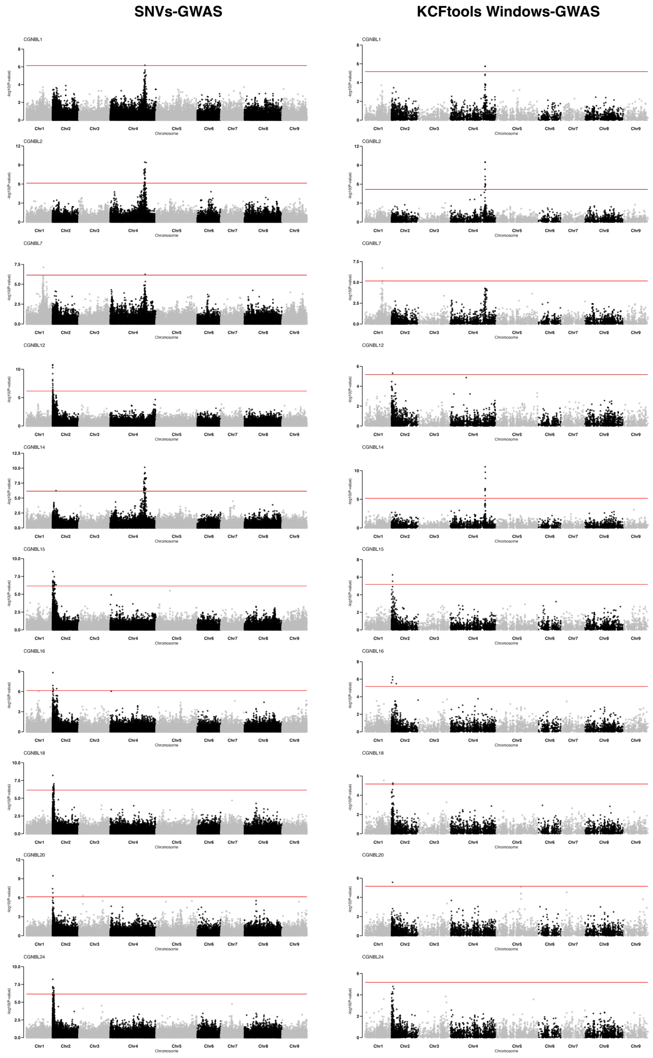

**Supplementary Figure S8:** Manhattan plots of Genome-Wide Association Studies (GWAS) of bremia resistance traits using the KCFtools windows as genotype data (right) and SNVs as genotype data(left)

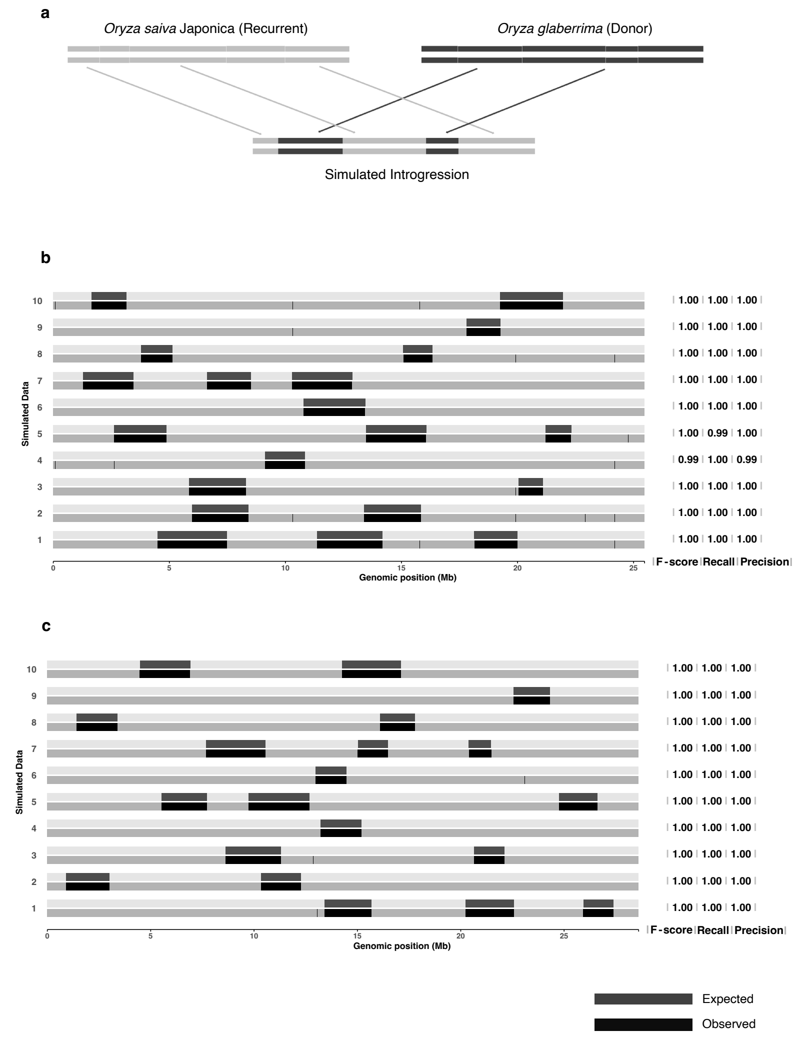

**Supplementary Figure S9:** Overview of the simulation experiment and results. a. figure shows the representative workflow of introducing the introgression from *Oryza glaberrima* to *Oryza sativa*. b. Results showing the expected (top bands) and observed (bands) introgressions from the ten simulated datasets while choosing *O. glaberrima* (donor parent) as reference. The benchmarking scores are mentioned on the right for each iterations. b. expected and observed variable regions on the simulated data, while using *O. sativa* (recurrent parent) as reference

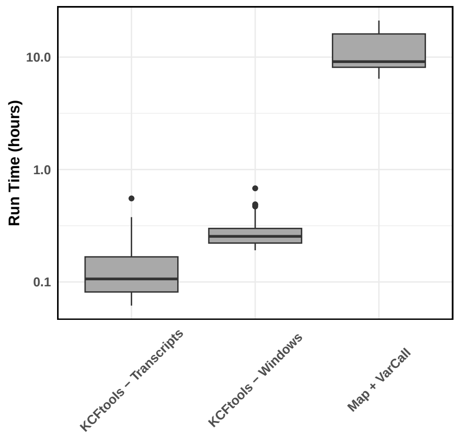

**Supplementary Figure S10:** Runtime comparison between KCFtools and a conventional variant-calling workflow. Boxplots summarize runtimes for 198 lettuce whole genome resequencing (WGRS) datasets queried against the *L. sativa* reference genome. Two KCFtools configurations were tested: using annotated transcripts as windows ("KCFtools – Transcripts") and using 50kb non-overlapping genomic segments as windows ("KCFtools – Windows"). The conventional workflow ("Map + VarCall") consisted of read mapping with BWA-MEM, BAM merging and indexing with SAMtools, and variant calling with DeepVariant. All analyses were performed using 10 threads. Reported KCFtools runtimes include the $k$-mer counting step performed with KMC.
